# Separate roles of LMAN1 and MCFD2 in ER-to-Golgi trafficking of factor V and factor VIII

**DOI:** 10.1101/2022.09.25.509417

**Authors:** Yuan Zhang, Zhigang Liu, Bin Zhang

## Abstract

Mutations in *LMAN1* and *MCFD2* cause the combined deficiency of FV and FVIII (F5F8D). LMAN1 and MCFD2 form a protein complex that transport FV and FVIII from the endoplasmic reticulum to the Golgi. Although both proteins are required for the cargo receptor function, little is known about specific roles of LMAN1 and MCFD2 in transporting FV/FVIII. We used different LMAN1 and MCFD2 deficient cell lines to investigate the LMAN1/MCFD2-dependent FV/FVIII secretion pathway. LMAN1 deficiency led to more profound decreases in FV/FVIII secretion in HEK293T and HepG2 cells than in HCT116 cells, suggesting regulation of cargo transport by the LMAN1/MCFD2 pathway varies in different cell types. Using these cell lines, we developed functional assays to accurately assess pathogenicity of recently reported potential LMAN1 and MCFD2 missense mutations. LMAN1 with mutations abolishing carbohydrate binding can still partially rescue FV/FVIII secretion, suggesting that N-glycan binding is not absolutely required for FV/FVIII transport. Surprisingly, overexpression of either WT or mutant MCFD2 is sufficient to rescue FV/FVIII secretion defects in LMAN1 deficient cells. These results suggest that cargo binding and transport are carried out by MCFD2 and that LMAN1 primarily serves as a shuttling carrier of MCFD2. Finally, overexpression of both LMAN1 and MCFD2 does not further increase FV/FVIII secretion, suggesting that the amount of the LMAN1-MCFD2 receptor complex is not a rate-limiting factor in ER-Golgi transport of FV/FVIII. This study provides new insight into the molecular mechanism of F5F8D and intracellular trafficking of FV and FVIII.

**Key Points:**

- Efficient ER-to-Golgi transport of FV and FVIII requires the LMAN1-MCFD2 cargo receptor complex.
- MCFD2 functions as a primary interacting partner of FV/FVIII cargo and LMAN1 primarily serves as a shuttling carrier of MCFD2.

## Introduction

Combined deficiency of coagulation factor V (FV) and factor VIII (FVIII) (F5F8D) is characterized by the simultaneous decreases of FV and FVIII antigen and activity levels in plasma to 5–30% of normal (1, 2). As a rare autosomal recessive disorder, F5F8D is often associated with consanguineous marriages, with the highest estimated occurrence at 1:100,000 among Middle Eastern Jews and non-Jewish Iranians (3–5). Patients with F5F8D exhibit mild-to-moderate bleeding symptoms. Approximately 70% of F5F8D cases are attributable to *LMAN1* (lectin, mannose binding 1) mutations (6), and 30% to *MCFD2* (multiple coagulation factor deficiency protein 2) mutations (1, 7, 8). LMAN1, which is also called ERGIC-53, is a 53 kD homo-hexameric transmembrane protein, and belongs to the family of L-type animal lectins (9). MCFD2 is a 16-kDa soluble monomeric protein with two Ca^2+^-binding motifs known as EF-hand domains. LMAN1 and MCFD2 form a Ca^2+^-dependent complex with 1:1 stoichiometry, and cycles between the ER and the ER–Golgi intermediate compartment (ERGIC) (7, 10). The LMAN1-MCFD2 complex serves as a cargo receptor for FV and FVIII, and facilitates their transportation from the ER to the Golgi (7, 10–12). Besides FV and FVIII, potential cargo for this receptor complex include α1-antitrypsin (AAT) and other proteins (11–19).

The mechanism of how the LMAN1-MCFD2 complex transports its cargo is not clear. LMAN1 has all the features of a cargo receptor: it is a type-1 transmembrane protein that has a carbohydrate binding domain (CRD) in the ER lumen and a short cytoplasmic domain that contains ER exit and retrieval motifs (7, 10, 20). However, LMAN1 cannot function as a transport receptor for FV, FVIII, AAT and perhaps other cargo in the absence of MCFD2, as MCFD2 deficiency leads to the same or more severe symptoms as LMAN1 deficiency (1, 12). The requirement of a soluble cofactor MCFD2 in the LMAN1-MCFD2 cargo receptor complex has been an enigma, suggesting a more complex trafficking mechanism than previously characterized cargo receptors in yeast. The CRD of LMAN1 contains separable binding sites for MCFD2 and mannose (20). Both LMAN1 and MCFD2 were reported to interact with cargo (10, 20, 21). Mannose binding activity of LMAN1 is presumably important for cargo selection, but it is not directly demonstrated. A LMAN1-binding deficient MCFD2 missense mutant can still bind to FVIII, suggesting that MCFD2 interaction with FVIII is independent of LMAN1 (10). The EF-hand domains of MCFD2 not only bind to LMAN1 but also interact with FV and FVIII (21). LMAN1 was recently shown to interact with AAT and this interaction is independent of MCFD2 (22).

LMAN1 and MCFD2 deficiencies in mice also lead to decreased plasma FV, FVIII and AAT levels (11, 12). Mouse studies also revealed a strain-specific partial lethal phenotype in LMAN1 deficient mice, but not in MCFD2-deficient mice, suggesting distinct functions of the two proteins other than transporting FV/FVIII (11, 12). To further understand the molecular mechanism of F5F8D and the regulation of ER-Golgi trafficking of FV and FVIII, we developed a complementation assay in *LMAN1* and *MCFD2* knockout (KO) cells to rapidly test functions of LMAN1 and MCFD2 variants, as well as features of FV/FVIII required for receptor-mediated secretion. We demonstrate that reduction of FV/FVIII secretion varied greatly among different KO cells, suggesting that regulation of cargo transport by the LMAN1/MCFD2 pathway varies in different cell types. We provide evidence that carbohydrate binding is not essential for the FV/FVIII transport function of LMAN1. Surprisingly, overexpression of MCFD2 in *LMAN1* KO cells is sufficient to recue FV/FVIII secretion defects. These results suggest that cargo binding and transport are carried out by MCFD2 and that LMAN1 primarily serves as a carrier of MCFD2.

## Methods

### Cells

HepG2 cells were grown in ATCC-formulated Eagle’s Minimum Essential Medium (EMEM) supplemented with 10% FBS, 100 IU/ml penicillin and 100 IU/ml streptomycin at 37°C and in 5% CO2. Human embryonic kidney 293T cells were grown in Dulbecco’s Modified Eagle Medium (DMEM) supplemented with 10% FBS, 100 IU/ml penicillin and 100 IU/ml streptomycin at 37°C and in 5% CO2. HCT116 cells were grown in McCoy’s 5A Medium supplemented with 10% FBS, 100 IU/ml penicillin and 100 IU/ml streptomycin at 37°C and in 5% CO2. Cells were transfected using FuGENE 6 (Promega, Madison, WI) or lipofectamine 3000 (Thermo Fisher Scientific, MA) according to the manufacturer’s instructions. Generation of cell lines stably expressing LMAN1 and MCFD2 was carried out as previously described (22).

### Plasmid construction

Construction of the mutant constructs N156A, H178A, W67S, Δβ1 of LMAN1 and D129E of MCFD2 were described previously (7, 10, 20). Missense mutations (V147I and V100D) were introduced into the plasmid pED-FLAG-LMAN1 and pcDNA-MCFD2-Myc separately using the QuickChange site-directed mutagenesis II XL kit from Agilent (Santa Clara, CA). FVIII mutant constructs Δ807–816 (deletion of amino acids 807-816 of FVIII) and mut 807-816 (replacement of all amino acids between 807 and 816 of FVIII with alanine) were prepared by polymerase chain reaction (PCR) using the QuickChange site-directed mutagenesis II XL kit. The pMT2-FVIII-WT construct encoding full-length FVIII (23) was used as the template for PCR. Plasmid pED-FV encoding full-length FV was a gift from Dr. Rodney M. Camire (Children’s Hospital of Pennsylvania). All mutant constructs were confirmed by Sanger sequencing for the presence of desired mutations and the absence of unintended mutations.

### Reagents

Rabbit polyclonal antibody and mouse monoclonal antibody against FLAG were purchased from Sigma-Aldrich (St. Louis, MO). Monoclonal anti-Myc antibody was purchased from Santa Cruz Biotechnology (Santa Cruz, CA). Mouse monoclonal anti-AAT antibody was purchased from Proteintech (Rosemont, IL). Rabbit monoclonal anti-LMAN1 antibody was purchased from Abcam (Cambridge, MA). Protein A/G Plus-agarose beads were purchased from Santa Cruz Biotechnology. D-Mannose agarose was purchased from Sigma–Aldrich.

### Establishment of LMAN1-deficient HCT116 cells

The HCT116 cell line is derived from colorectal cancer with microsatellite instability. The exon 8 of *LMAN1* contains a microsatellite site with a string of 9 adenosines (9A). HCT116 cells consist of mostly heterozygous population with alleles of 8A and 9A (8A/9A) at this site (24). We discovered that a minority of HCT116 cells have homozygous 8A alleles (8A/8A). Colonies derived from single cells were genotyped to identify cells with homozygous 8A/8A and heterozygous 8A/9A alleles. The lack of LMAN1 expression in homozygous 8A/8A KO cells was confirmed by immunoblotting.

### FVIII activity and FVIII/FV antigen analysis

FVIII activity was measured by a chromogenic assay using the Coatest SP4 FVIII kit from DiaPharma (West Chester, OH). FVIII antigen was quantified by ELISA using the VisuLize FVIII antigen kit from Affinity Biologicals (Ancaster, ON, Canada). FV antigen was quantified by ELISA using matched-pair antibody set for human FV antigen from Affinity Biologicals. All assays were performed according to manufacturers’ instructions.

### Immunoprecipitation and mannose binding assay

Immunoprecipitation of LMAN1 and MCFD2 was performed as previously described (10). Mannose binding assay was performed as previously described (20), with some modifications. Briefly, 293T cells were harvested on ice in homogenate buffer 48 h after transfection with LMAN1 expression constructs. Cells were homogenized and cleared by centrifugation at 500 g for 10 min. Membrane fraction from the postnuclear supernatant was pelleted at 100,000g for 1 hour. The pellet was solubilized for 1 h in lysis buffer (10mM Tris-HCl, pH 7.4, 150mM NaCl, 10mM CaCl_2_, 1mM MgCl_2_ containing 1% Triton X-100, and protease inhibitors), followed by a centrifugation at 100,000g for 1 h and dialysis of the supernatant overnight against binding buffer (10mM Tris-HCl, pH 7.4, 150mM NaCl, 10mM CaCl_2_, 1 mM MgCl_2_, and 0.15% Triton X-100). The dialysate was incubated with D-mannose agarose beads overnight, and bound LMAN1 was eluted using 0.2 M mannose in binding buffer. Eluted LMAN1 was detected by immunoblotting analysis using a rabbit anti-FLAG antibody.

### Statistical analysis

FVIII activity, FVIII antigen and FV antigen assay results were analyzed using the Student’s t test for comparison between 2 groups and by one-way analysis of variance (ANOVA) for comparisons of >2 groups. P-values <0.05 were considered significant for all assays.

### Data Sharing Statement

For original data, please contact.

## Results

### Extent of decreases in FV and FVIII secretion varies in different *LMAN1* and *MCFD2* KO cells

Using the CRISPR-Cas9 technology, we previously established 293T and HepG2 cells with *LMAN1* or *MCFD2* KO and found reduced rates of ER-Golgi transport of AAT in these cells (22). We have now also established LMAN1-deficient HCT116 (HCT116^-/-^) cells with natural frameshift mutations in exon 8 of *LMAN1* (Fig. S1) (24). LMAN1 and MCFD2 are expressed in HepG2 cells at much higher levels than in 293T and HCT116 cells (Fig. 1A). As expected, *LMAN1* KO cells have marked reduction in MCFD2 levels due to its reliance on LMAN1 for intracellular retention, whereas *MCFD2* KO had no effects on LMAN1 levels (Fig. 1A). Secretion of endogenous FV was analyzed by measuring FV antigen levels in conditioned media of WT and KO HepG2 cells. Secretion of FV was also analyzed in WT and KO 293T or HCT116 cells by measuring FV antigen levels in conditioned media of cells transfected with a FV expression plasmid. Results showed that FV secretion levels were reduced to ~50% of the WT level in both *LMAN1* and *MCFD2* KO 293T and HepG2 cell lines (Fig. 1B). However, FV secretion in HCT116^-/-^ cells only decreased mildly, to ~80% of the HCT116^+/-^ level (Fig. 1B). To measure secretion of FVIII, a FVIII expression plasmid was transfected into WT and KO cells, and FVIII activity and antigen levels in conditioned media were detected 48 h post transfection. FVIII activity and antigen levels decreased in all 293T and HepG2 KO cell lines (293T^LMAN1-^, 293T^MCFD2-^, HepG2^LMAN1-^, HepG2^MCFD2-^) to less than 10% level of WT cells (Fig. 1C). However, reduction of FVIII in conditioned media of HCT116^-/-^ cells was less profound, to only ~50% level of HCT116^+/-^ cells (Fig. 1C). Moderate decrease in FVIII secretion in HCT116^-/-^ cells is consistent with a previous report (25).

**Figure 1.**
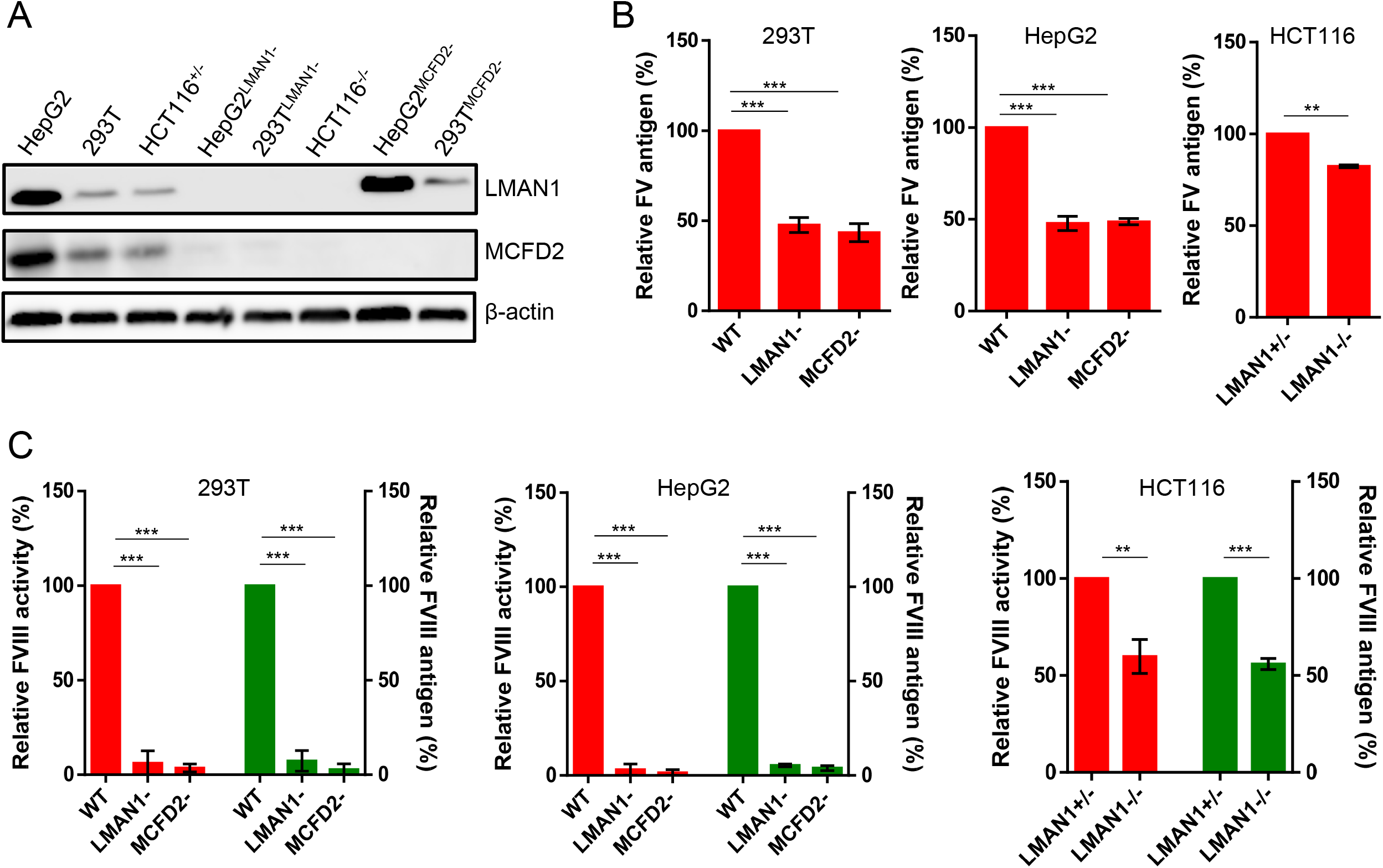
Secretion levels of FVIII and FV in different cell lines. (A) LMAN1 and MCFD2 levels in WT, HepG2^LMAN1-^, HepG2^MCFD2-^, 293T^LMAN1-^ and 293T^MCFD2-^ cells, as well as HCT116^+/-^ and HCT116^-/-^ cells. (B) Conditioned media were collected from HepG2 cell lines 48 h after fresh medium change. A FV expression construct was transfected into the indicated 293T and HCT116 cell lines and conditioned media were collected 48 h post transfection. FV levels were measured by ELISA and plotted as percentages of WT cell levels. (C) A FVIII expression construct was transfected into the indicated cell lines and conditioned media were collected 48 h post transfection. FVIII activity and antigen levels in conditioned media were measured and plotted as percentages of WT cell levels. Data presented are means of 3 independent experiments, and the error bars represent standard deviations. **P<0.01, ***P<0.001.

### The SDLLMLLRQS sequence in FVIII B domain is not required for LMAN1/MCFD2-dependent secretion

A recent study reported a putative MCFD2-binding segment from the B domain of FVIII (SDLLMLLRQS at residues 807-816) (25). To test the role of this sequence in the LMAN1-MCFD2 secretion pathway of FVIII, we constructed FVIII with the deletion of these residues [Δ(807-816)] or the alanine replacement of all residues in this segment [mut(807-816)]. Secretion of WT and mutant FVIII was compared in WT, *LMAN1* KO and *MCFD2* KO cell lines. Deletion or mutation of this segment did not cause reduction of activity and antigen levels of FVIII secreted in WT 293T, HepG2 and HCT116 cell lines (Fig. S2). FVIII activity and antigen levels of both Δ(807-816) and mut(807-816) mutants were reduced in conditioned media of both *LMAN1* KO and *MCFD2* KO 293T and HepG2 cells, as well as in LMAN1-deficient HCT116^-/-^ cells, to levels identical to WT FVIII (Fig. S2). These results suggest that the SDLLMLLRQS sequence in the B domain is not required for LMAN1/MCFD2-dependent secretion of FVIII.

### LMAN1 with mutations abolishing carbohydrate binding can still partially rescue FVIII secretion

To ask whether re-expression of LMAN1 in *LMAN1* KO cells can rescue the FVIII secretion defects, we co-transfected FVIII and different LMAN1 expression constructs into 293T^LMAN1-^ cells (Fig. 2A). Only functional LMAN1 or MCFD2 molecules are expected to rescue FVIII secretion defects in KO cells. As expected, co-transfection of WT LMAN1 can fully restore active FVIII secretion (Fig. 2B-C). In contrast, no rescue of FVIII secretion occurred with co-transfection of the W67S and Δβ1 variants (Fig. 2B-C). W67S is a patient mutation that abolishes both MCFD2 binding and mannose binding (20). Δβ1 mutant has a deletion of the first β-sheet in the CRD of LMAN1, which is directly involved in MCFD2 binding, thus abolishing MCFD2 binding without affecting mannose binding (11). V147I is a recently reported missense variant in LMAN1 found in a F5F8D patient from China (26). However, co-transfection of the V147I variant resulted in FVIII secretion at a level comparable to the co-transfection of WT LMAN1 (Fig. 2A-C), indicating that this variant did not affect the LMAN1 function in FVIII secretion.

**Figure 2.**
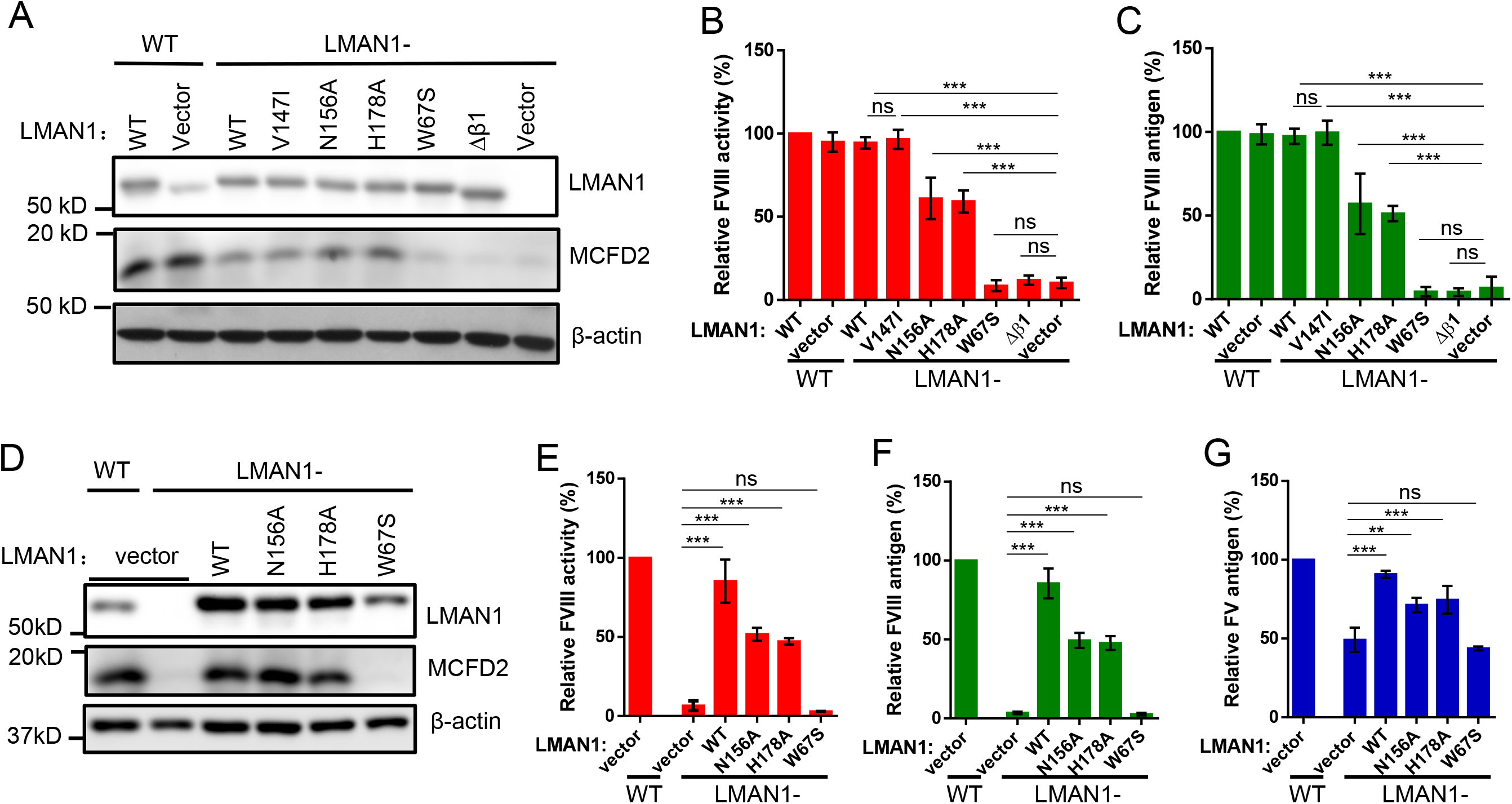
Rescue of FV and FVIII secretion in *LMAN1* KO cells by LMAN1 variants. (A) WT 293T cells were transfected with WT LMAN1 or the vector. 293T^LMAN1-^ cells were transfected with WT LMAN1, the indicated LMAN1 variants or the vector. Forty-eight hours post transfection, conditioned media were collected and cell lysates were subjected to immunoblotting analyses with anti-LMAN1, anti-MCFD2 and anti-β actin antibodies. FVIII activity (B) and antigen (C) levels in conditioned media were measured and plotted as percentages of WT cell levels. (D) WT LMAN1 and the indicated LMAN1 variants were stably expressed into HepG2^LMAN1-^ cells and LMAN1 expression levels were compared with vector-transduced WT HepG2 and HepG2^LMAN1-^ cells by immunoblotting. (E-F) A FVIII expression construct was transfected into HepG2^LMAN1-^ cells stably expressing indicated LMAN1 variants and conditioned media were collected 48 h post transfection. FVIII activity and antigen levels in conditioned media were measured and plotted as percentages of WT cell levels. (G) Conditioned media were collected 48 h after fresh medium change. FV antigen levels were measured by ELISA and plotted as percentages of WT cell levels. FV and FVIII level data presented in this figure are means of 3 independent experiments, and the error bars represent standard deviations. **P<0.01, ***P<0.001. ns, not significant.

Using this LMAN1 complementation assay, we found that LMAN1 variants with point mutations in the mannose-binding site (N156A and H178A) can still rescue FVIII secretion to ~60% level of WT LMAN1 (Fig. 2B-C) in 293T cells. To rule out the possibility that the surprising rescue of FVIII secretion by N156A and H178A mutants were an artifact of overexpression, we decreased LMAN1 expression to levels comparable or lower than that in WT cells. Under these conditions, N156A and H178A mutants can still rescue most of FVIII secretion (Fig. S3). Next, we established HepG2^LMAN1-^ cell lines that stably expressing WT, N156A, H178A and W67S variants of LMAN1 using retroviral expression vectors (Fig. 2D). In this system, transfected FVIII secretion was also partially rescued in HepG2^LMAN1-^ cell lines stably expressing N156A and H178A variants (Fig. 2E-F). In addition, WT LMAN1 rescued endogenous FV secretion in HepG2^LMAN1-^ cells to ~90% of WT cells, whereas N156A and H178A variants rescued FV secretion to ~70% level of WT cells (Fig. 2G).

The N156A mutation changes a critical amino acid in the carbohydrate binding pocket and was shown to abolish mannose binding without affecting MCFD2 binding (9). We have previously shown by isothermal titration calorimetry that the H178A mutation abolished dimannose binding without affecting Ca^2+^ binding (27). Here, we directly confirmed mannose binding deficiency of both N156A and H178A mutants in a mannose binding assay (Fig. 3A). On the other hand, mannose binding activity is maintained in the V147I variant (Fig. 3A). To test the LMAN1-MCFD2 interaction, we performed a co-immunoprecipitation (co-IP) assay. We included WT as a positive control, and W67S and Δβ1 as negative controls. Results showed that the V147I, N156A and H178A variants could still co-immunoprecipitate with MCFD2 (Fig. 3B), suggesting that the LMAN1-MCFD2 interaction was not disrupted by these mutations. Taken together, these results suggest that carbohydrate binding is not absolutely required for the cargo receptor function of LMAN1, and that V147I is not a deleterious mutation.

**Figure 3.**
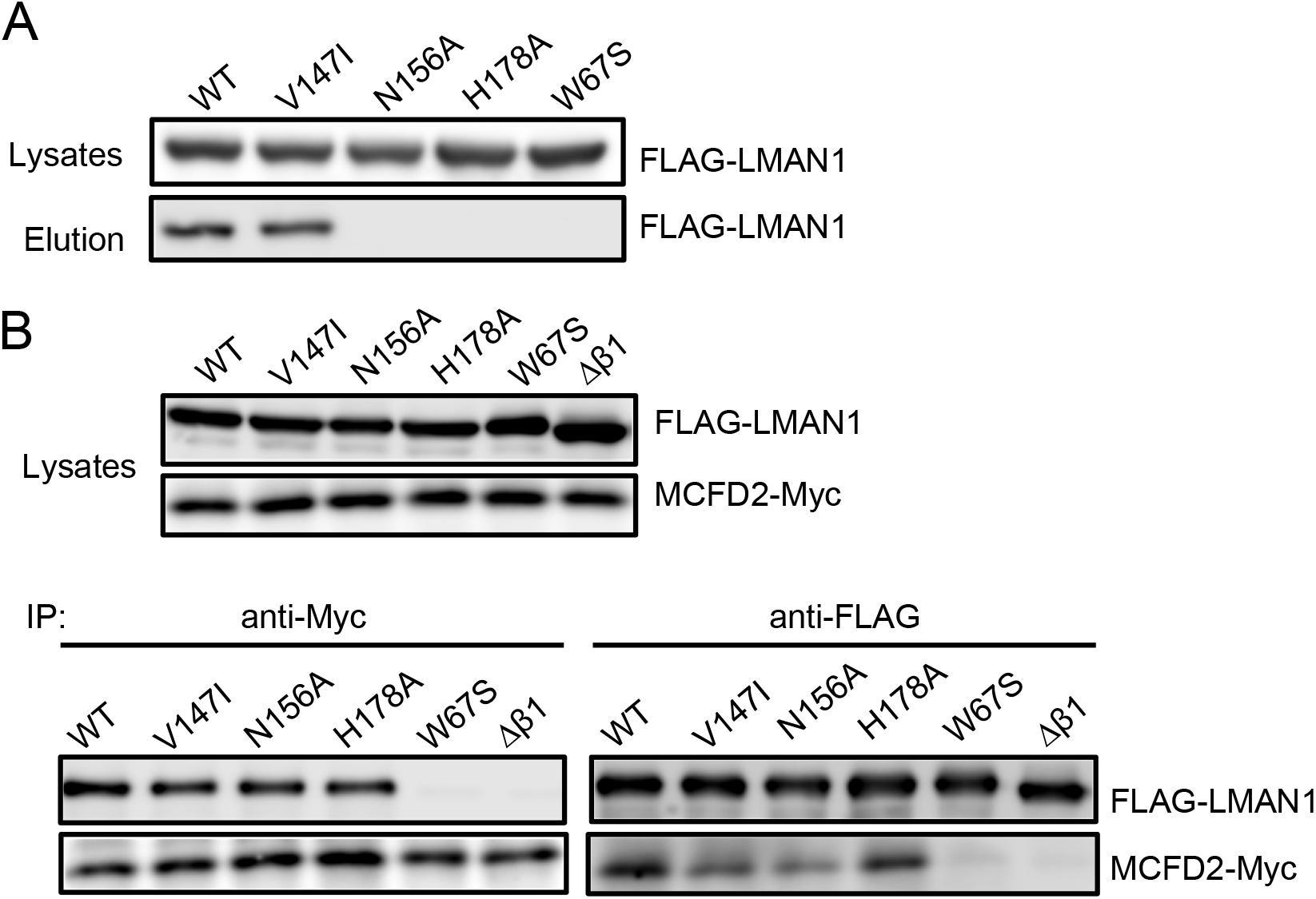
Interactions of LMAN1 variants with mannose and MCFD2. (A) Mannose binding assay. 293T cells were transfected with the FLAG-tagged WT and the indicated LMAN1 variants. Cell lysates were loaded onto a mannose agarose column. The bound LMAN1 was eluted from the column and detected by immunoblotting. (B) Co-IP of LMAN1 variants and MCFD2. 293T cells were co-transfected with FLAG-tagged WT and LMAN1 mutants and a Myc-tagged MCFD2. Cell lysates were immunoprecipitated with anti-Myc for MCFD2 and anti-FLAG for LMAN1 and detected by immunoblotting.

### The V100D variant of MCFD2 is a hypomorphic mutation

To test MCFD2 mutant functions, we created HepG2 and 293T cell lines that stably expressing WT and different mutant MCFD2 by transducing cells with retroviruses carrying MCFD2 expression constructs into 293T^LMAN1-^ and HepG2^LMAN1-^ cells (Fig. 4A, 4E). These cells were then transfected with a FVIII expression construct to assess the secretion of FVIII. WT MCFD2 restored FVIII secretion in both HepG2^MCFD2-^ (Fig. 4B-C) and 293T^MCFD2-^ cells (Fig. 4F-G) to ~80% of WT cells. Expression of WT MCFD2 restored endogenous FV secretion in HepG2^MCFD2-^ cells and transfected FV secretion in 293T^MCFD2-^ cells to ~70-80% of WT cells (Fig. 4D, 4H). D129E and Y135N are patient mutations localized to the second EF-hand domain and causes the disease due to disruption of the LMAN1-MCFD2 complex (21). We observed small but significant increases FVIII secretion in both HepG2^MCFD2-^ and 293T^MCFD2-^ cells that expressed D129E and Y135N mutants compared to the vector control (Fig. 4B-C and Fig. 4F-G). Expression of D129E and Y135N mutants did not significantly increase FV secretion (Fig. 4D, 4H), likely due to higher basal levels of FV secretion in *MCFD2* KO cells. Using this MCFD2 complementation assay, we tested the function of the V100D variant of MCFD2. V100D is a variant reported in a Tunisian patient, and is localized at the helix 2 of the first EF hand domain (28). Results showed that the V100D variant was also able to partially rescued FVIII secretion to ~50% level of WT cells in HepG2^MCFD2-^ cells (Fig. 4B-C) and > 80% in 293T^MCFD2-^ cells (Fig. 4F-G). Similarly, this variant also partially rescued the endogenous FV secretion in both cell lines (Fig. 4D, 4H). The V100D variant could also co-immunoprecipitate with LMAN1, indicating that it can form a complex with LMAN1 (Fig. S4). These results suggest that the V100D mutation is hypomorphic and likely not a disease-causing mutation.

**Figure 4.**
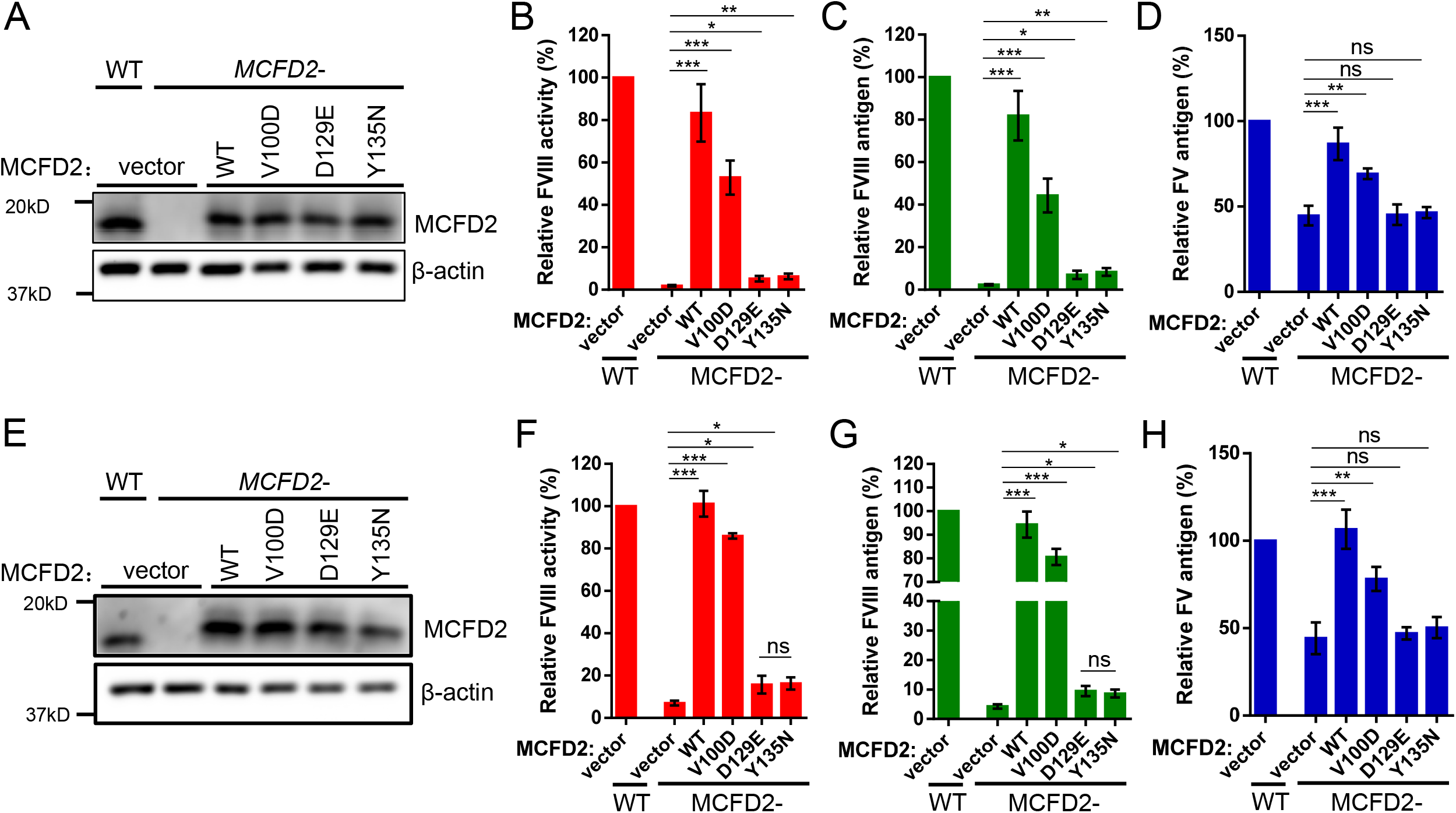
Rescue of FV and FVIII secretion in *MCFD2* KO cells by MCFD2 variants. (A) WT MCFD2 and the indicated MCFD2 variants were stably expressed in HepG2^MCFD2-^ cells and MCFD2 expression levels were compared with vector-transduced WT HepG2 and HepG2^MCFD2-^ cells by immunoblotting. A FVIII expression construct was transfected into these cells and conditioned media were collected 48 h post transfection. FVIII activity (B) and antigen (C) levels in conditioned media were measured and plotted as percentages of WT cell levels. (D) Conditioned media were collected 48 h after fresh medium change. FV antigen levels were measured by ELISA and plotted as percentages of WT cell levels. (E) WT MCFD2 and the indicated MCFD2 variants were stably expressed in 293T^MCFD2-^ cells and MCFD2 expression levels were compared with vector-transduced WT 293T and 293T^MCFD2-^ cells by immunoblotting. A FVIII expression construct or a FV expression construct was transfected into these cells and conditioned media were collected 48 h post transfection. FVIII activity (F), FVIII antigen (G) and FV antigen (H) levels in conditioned media were measured and plotted as percentages of WT cell levels. FV and FVIII level data presented in this figure are means of 3 independent experiments, and the error bars represent standard deviations. *P<0.05, **P<0.01, ***P<0.001. ns, not significant.

### Overexpression of MCFD2 in *LMAN1* KO 293T cells restores FVIII secretion

As noted above, 293T^MCFD2-^ and HepG2^MCFD2-^ cells stably expressing D129E or Y135N mutants had FVIII secretion levels ~2-fold higher than the vector control (Fig. 4B-C, 4F-G). One possible reason for this could be that these mutants may retain residual LMAN1-binding activities (8), and a trace amount of the LMAN1-MCFD2 complex is responsible for the increased FVIII secretion. To exclude this possibility, we established stable cell lines that express WT MCFD2 and different MCFD2 variants in 293T^LMAN1-^ cells. Steady-state levels of WT and variant MCFD2 were similar in cell lysates (Fig. S5). Interestingly, FVIII secretion still increased by 2-fold in cells expressing either WT or MCFD2 variants compared to 293T^LMAN1-^ cells with vector control after transfection of a FVIII expression construct (Fig. S5), suggesting that the increased FVIII secretion was due to the expression of MCFD2 or its variants, not the formation of a trace amount of the LMAN1-MCFD2 complex.

To test whether overexpression of MCFD2 can further rescue FVIII secretion, we co-transfected MCFD2 and FVIII expression constructs into 293T^MCFD2-^ and 293T^LMAN1-^ cells and measured antigen and activity levels of secreted FVIII. MCFD2 was transfected in two doses to control the amounts of protein expression. Protein levels of transfected MCFD2 in both KO cells were 5-6 fold higher than the endogenous MCFD2 level in WT cells with 50 ng plasmid DNA, and 2-3 fold higher than WT cells with 20 ng plasmid DNA (Fig. 5A, 5D). In both cell lines, all MCFD2 variants (WT, V100D, D129E and Y135N) rescued FVIII secretion. Higher MCFD2 expression rescued FVIII secretion to ~90% levels of WT MCFD2, while lower MCFD2 expression led to 50-70% rescue (Fig. 5C, 5F). On the other hand, overexpression of LMAN1 in 293T^MCFD2-^ cells had no effect on FVIII secretion (Fig. S6). Overexpression of MCFD2 variants also led to near complete rescue of FV secretion in both cell lines (Fig. 5B, 5E). In addition, overexpression of MCFD2 in 293T^MCFD2-^ and 293T^LMAN1-^ cells also rescued the FVIIIΔ(807-816) secretion (Fig. S7), providing further evidence that this B-domain sequence is not important in LMAN1/MCFD2-dependent secretion of FVIII.

**Figure 5.**
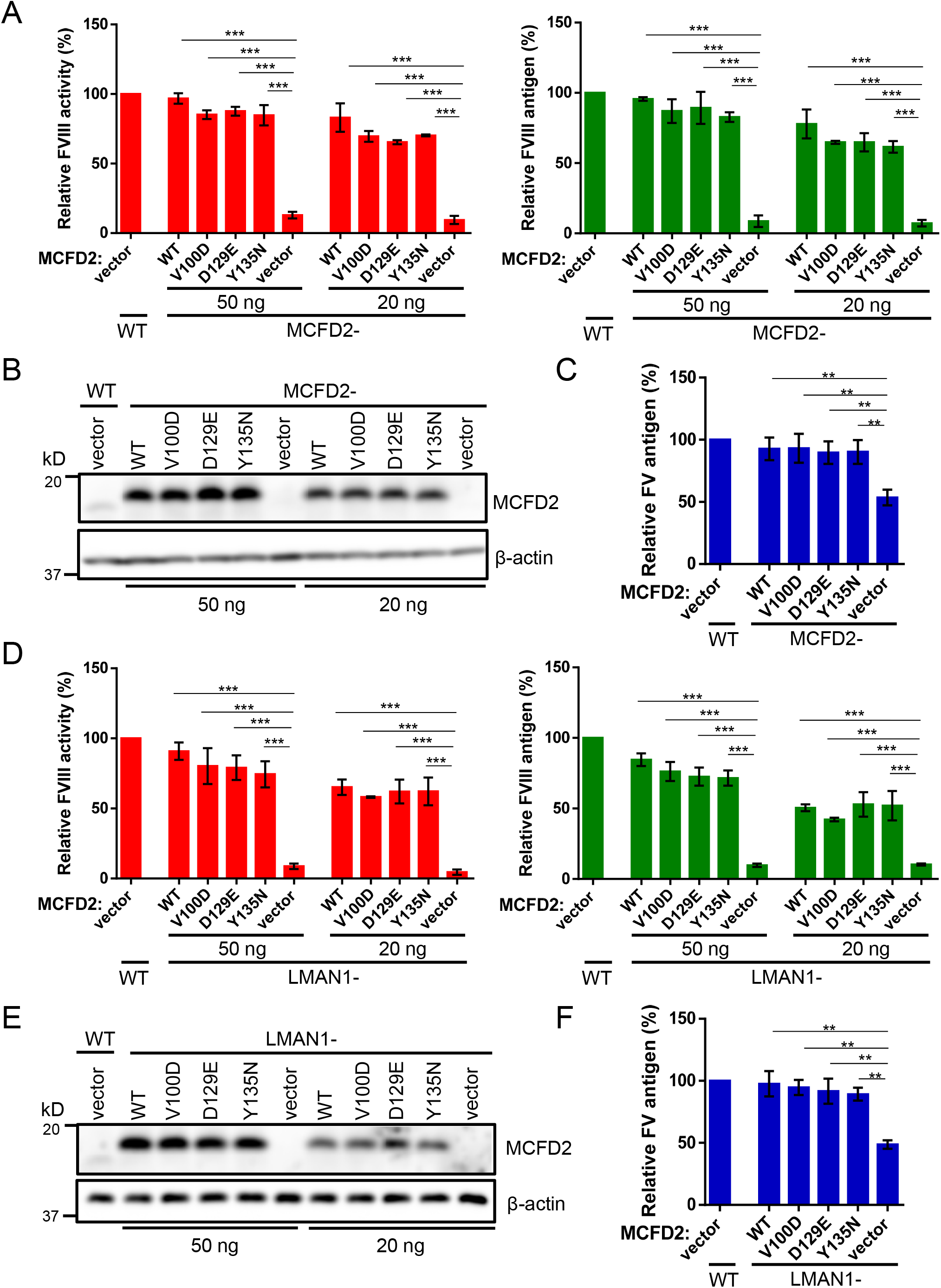
Overexpression of MCFD2 rescues FVIII secretion in LMAN1 KO and MCFD2 KO cells. (A) A FV or FVIII expression construct and vectors expressing the indicated MCFD2 variants were co-transfected into 293T^MCFD2-^ cells in 24 well plates in two doses (50 ng and 20 ng). MCFD2 expression levels in cell lysates were compared with vector-transfected WT 293T and 293T^MCFD2-^ cells by immunoblotting. (B) FV antigen levels in conditioned media were measured 48 h post transfection and plotted as percentages of WT cell levels. (C) FVIII activity and antigen levels in conditioned media were measured 48 h post transfection and plotted as percentages of WT cell levels. (D) A FV or FVIII expression construct and vectors expressing the indicated MCFD2 variants were co-transfected into 293T^LMAN1-^ cells in 24 well plates in two doses (50 ng and 20 ng). MCFD2 expression levels in cell lysates were compared with vector-transfected WT 293T and 293T^MCFD2-^ cells by immunoblotting. MCFD2 expression levels in cell lysates were compared with vector-transfected WT 293T and 293T^LMAN1-^ cells by immunoblotting. (E) FV antigen levels in conditioned media of were measured 48 h post transfection and plotted as percentages of WT cell levels. (F) FVIII activity and antigen levels in conditioned media were measured 48 h post transfection and plotted as percentages of WT cell levels. FVIII activity and antigen data presented are means of 3 independent experiments, and the error bars represent standard deviations. **P<0.01, ***P<0.001.

### Overexpression of the LMAN1-MCFD2 complex does not further increase FV and FVIII secretion

Previous results showed that LMAN1 and MCFD2 form a complex with 1:1 stoichiometry (10). However, overexpression of LMAN1 did not increase endogenous MCFD2 beyond the WT cell level (Fig. 2). Overexpression of MCFD2 also did not increase the endogenous LMAN1 level (10). These results suggest that levels of LMAN1 and MCFD2 are independently regulated. To increase LMAN1 and MCFD2 levels simultaneously in HepG2 cells, we transfected MCFD2 into HepG2^LMAN1-^ cells stably overexpressing LMAN1. The resulting cells express both LMAN1 and MCFD2 to 2-3 fold levels of WT HepG2 cells (Fig. 6A, lane 2). However, secreted endogenous FV levels in conditioned media of these cells were not significantly different from WT cells and cells overexpressing MCFD2 or LMAN1 alone, and higher than *LMAN1* KO cells (Fig. 6B). To assess whether increasing both LMAN1 and MCFD2 can increase FVIII secretion, we co-transfected LMAN1 and FVIII into 293T cells stably overexpressing WT MCFD2, which led to overexpression of both LMAN1 and MCFD2 (Fig. 6C, lane 2). FVIII levels in conditioned media of this cell line were unchanged compared to WT 293T cells or 293T cells overexpressing LMAN1 or MCFD2 alone (Fig. 6D). These results indicate that increasing the amount of the LMAN1-MCFD2 complex does not lead to increases in FV and FVIII secretion.

**Figure 6.**
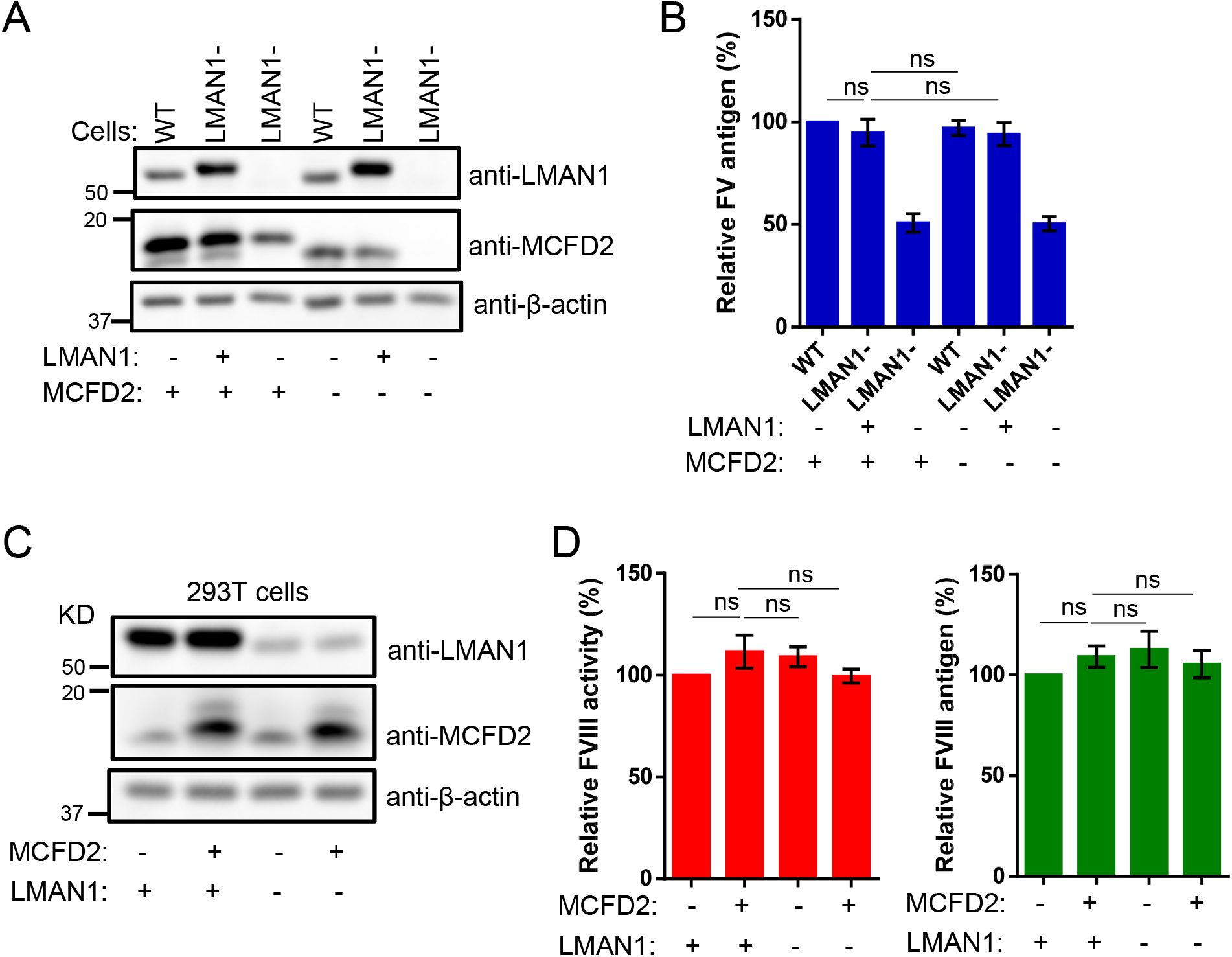
Overexpression of the LMAN1-MCFD2 complex does not further increase FV and FVIII secretion. (A) WT HepG2 and HepG2^LMAN1-^ cells with (+) or without (-) stably expressing LMAN1 were transfected with a MCFD2 expressing construct (+) or with an empty vector (-). Cell lysates were subjected to immunoblotting 48 h post transfection with the indicated antibodies. (B) FV antigen levels were measured in conditioned media 48 h post transfection and plotted as percentages of WT cell levels. (C) WT 293T cells with (+) or without (-) stably overexpressing MCFD2 were co-transfected with a FVIII expression construct and a LMAN1 expressing construct (+) or an empty vector (-) as indicated. Cell lysates were subjected to immunoblotting 48 h post transfection with the indicated antibodies. (D) FVIII activity and antigen levels were measured in conditioned media 48 h post transfection and plotted as percentages of WT cell levels. FV and FVIII level data presented are means of 3 independent experiments, and the error bars represent standard deviations. ns, not significant.

## Discussion

LMAN1 deficiency in all three tested cell lines led to decreased FV and FVIII secretion. MCFD2 deficiency in 293T and HepG2 cells also led to similar decreases in FV and FVIII secretion. These results suggest that these cell lines can be used to model F5F8D *in vitro*. However, the extent of secretion decrease varies greatly in different cells, suggesting that regulation of cargo transport by the LMAN1/MCFD2 pathway varies in different cell types. In particular, FV/FVIII secretion in HCT116 cells appears less dependent on the LMAN1-MCFD2 complex, and suggests the existence of an alternative pathway for ER-Golgi transport of FV/FVIII or other cargo receptors with functional overlap. FVIII is primarily expressed in endothelial cells (29, 30). Human FV is expressed in hepatocytes and taken up by megakaryocytes via endocytosis (31). Although none of the cell lines naturally synthesize FVIII, HepG2 cells have endogenous FV expression and express much higher levels of LMAN1 and MCFD2. Curiously, the extent of decrease in FV secretion is much less than the decrease in FVIII secretion in the same KO cells. This is in contrast to both F5F8D patients (1) and LMAN1/MCFD2-deficient mice (12), which have moderate correlations in plasma FV and FVIII levels. This discrepancy may be due to intrinsic differences between FV and FVIII expressed *in vitro* (32).

Signals in FVIII required for the interaction with the LMAN1-MCFD2 complex have not been identified, although it is thought that N-glycans in cargo proteins play a major role in LMAN1 interaction. To our surprise, we found that the N156A and H178A mutations that abolish sugar binding were still able to rescue most of the FVIII secretion defects in *LMAN1* KO cells. This result may also explain why no missense mutations have been identified in the carbohydrate binding region of LMAN1 (1). Any LMAN1 missense mutation would have to severely affect the protein expression or function, as a hypomorphic mutation that reduces murine *Lman1* mRNA expression to 6-8% of WT level leads to intermediate plasma FV and FVIII reductions (33). There are only two LMAN1 missense mutations reported to date. The W67S mutation causes a MCFD2 binding defect in addition to the mannose binding defect (20, 34). The C558R mutation leads to an unstable protein (8). More missense mutations were identified in MCFD2, all of which abolish LMAN1 binding. These results suggest that LMAN1-MCFD2 complex formation, but not N-glycan binding of LMAN1, is absolutely required for the cargo receptor function. A possible explanation is that LMAN1 can interact with FVIII through a lectin-independent, direct protein-protein interaction. Our previous studies showed that cross-linking of FVIII with LMAN1 can still be observed in cells treated with N-glycosylation inhibitor tunicamycin (10). Targeting of procathepsin Z appears to require both an oligosaccharide chain and a surface-exposed peptide β-hairpin loop (35). MMP-9 N-glycosylation mutants also strongly co-IP with LMAN1 (19). Crystal structures of the LMAN1-CRD suggested potential protein-binding sites for cargo proteins (36, 37).

We also tested a recently reported MCFD2-binding motif from the FVIII B domain. It was reported that deletion of this putative motif caused reduced secretion of FVIII from HCT116 cells (25). However, we could not duplicate this result in HCT116 cells. FVIII is only moderately decreased in HCT116^-/-^ cells. We also noted that Yagi et al (25) cultured HCT116 cells in a different medium and used a codon-optimized version of FVIII construct, which could explain some differences in our results. No secretion defects of motif-deleted or mutated FVIII were observed in 293T and HepG2 cells either. Moreover, secretion of both WT FVIII and motif-deleted or motif-mutated FVIII was reduced to the same extent in all three *LMAN1* KO cell lines, as well as in 293T^MCFD2-^ and HepG2^MCFD2-^ cells. These results strongly suggest that this motif is either not a signal or not sufficient by itself to serve as a signal for LMAN1-MCFD2-dependent secretion of FVIII. Further studies are needed to identify signals recognized by the LMAN1-MCFD2 cargo receptor complex.

*LMAN1* and *MCFD2* KO cell lines provide convenient functional assays to accurately assess the pathogenicity of LMAN1 and MCFD2 mutations. The V147I variant is located at a highly conserved site and *in silico* analysis predicted that this is a pathogenic mutation (26). Our results showed that it can fully rescue FVIII secretion defects in LMAN1 KO cells, and has no detected defects in MCFD2 and mannose binding. Therefore it is likely not a disease-causing mutation. Reduction of FV and FVIII levels in this patient may result from a mutation in either *LMAN1* or *MCFD2* that was missed by DNA sequencing. There were previous reports of patients who were deficient in LMAN1 with no mutations identified in the exons and exon-intron junctions (1, 8). The impact of V100D variant on MCFD2 structure has been extensively studied with inconclusive results (28, 38–40). Circular dichroism analysis of the recombinant V100D protein suggested reduced structural stability of this variant (38). V100D is carried together in a heterozygous state with another missense mutation D81H in a homozygous state (28). We show here that this is a hypomorphic variant that is unlikely to cause F5F8D if inherited in a homozygous or compound heterozygous state.

Overexpression of both LMAN1 and MCFD2 rescued FV/FVIII secretion to levels comparable to, but not exceeding WT cells, suggesting that the LMAN1-MCFD2 cargo receptor level is not a rate limiting factor in ER-Golgi transport of FV/FVIII. All MCFD2 missense mutations identified to date are localized to the EF hand domains (21) that are important for LMAN1 binding (36, 37). Although proper cargo receptor function requires the LMAN1-MCFD2 complex formation, patients with *MCFD2* mutations generally have lower plasma FV/FVIII levels than patients with *LMAN1* mutations (1), suggesting a more direct role of MCFD2 in FV/FVIII transport. Here we present a surprising finding that MCFD2 alone could rescue FV/FVIII secretion defects in 293T^LMAN1-^ cells. Our previous studies showed that MCFD2 interacts with FV and FVIII independent of LMAN1, suggesting that MCFD2 contains distinct FV/FVIII and LMAN1 binding sites (21). LMAN1-deficient patients have only trace amounts of intracellular MCFD2 due to the requirement of LMAN1 for intracellular retention (7, 10). Our data suggest that LMAN1 and MCFD2 have distinct functions in cargo transport. MCFD2 is the “cargo capture module” that brings cargo proteins to the LMAN1-MCFD2 complex in the ER for packaging into COPII vesicles. The major function of LMAN1 in cargo transport is serving as an ER-Golgi shuttling carrier for MCFD2. The mannose binding site of LMAN1 may further stabilize the tertiary complex with FV/FVIII. Although LMAN1 could co-IP with AAT independent of MCFD2, this interaction is apparently not sufficient to form a stable transport-competent complex, as AAT secretion is also decreased in *MCFD2* KO cells (22). When overexpressed in *LMAN1* KO cells, the flux of MCFD2 protein may be sufficient to overcome the lack of LMAN1 carrier. This function of MCFD2 is clearly independent of LMAN1 binding as mutant MCFD2 that cannot bind LMAN1 can still rescue FVIII secretion in *LMAN1* KO cells. How does MCFD2 transport FV/FVIII without LMAN1? It could serve as a chaperonin protein that stabilizes cargo in the ER, or the MCFD2-cargo complex formation is sufficient to facilitate the ER exit of cargo by bulk flow.

In conclusion, our results demonstrate that LMAN1 and MCFD2 deficient cell lines provide valuable tools to study cargo receptor-mediated secretion of FV/FVIII and other proteins. Using these cells, we functionally characterized LMAN1 and MCFD2 variants from F5F8D patients and showed that regulation of cargo transport by the LMAN1/MCFD2 pathway varies in different cell types. We present evidence supporting a model in which MCFD2 functions as a primary interacting partner of FV/FVIII cargo and LMAN1 primarily serves as a vehicle that shuttles MCFD2 between the ER and the Golgi.

## ACKNOWLEDGEMENT

This study was supported by the National Institutes of Health (R01 HL094505 to B.Z.) and the Alpha-1 Foundation. Y.Z. is a recipient of the Judith Graham Pool Postgraduate Fellowship from the National Hemophilia Foundation.

## AUTHORSHIP

Contribution: Y.Z. and B.Z. designed the study; Y.Z. and Z.L. performed research; Y.Z., Z.L. and B.Z. analyzed data; Y.Z. and B.Z. wrote the manuscript, and Z.L. provided critical comments.

