## Supplemental figures for "Separate roles of LMAN1 and MCFD2 in ER-to-Golgi trafficking of factor V and factor VIII"

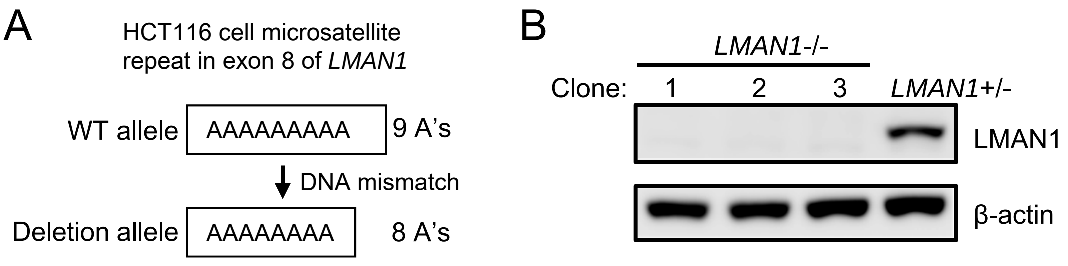


**Figure S1. Generation of LMAN1-deficient HCT116 cells**. (A) DNA mismatch repair deficiency leads to mostly heterozygous shortening of the microsatellite sequence in exon 8 of *LMAN1* in HCT116 cells. (B) Immunoblotting results show the lack of LMAN1 expression in clones 1, 2, and 3, which contain homozygous 8A/8A alleles at the microsatellite site. Similar reductions in FV/FVIII secretion were obtained from all three independent HCT116-/- cell clones, therefore, one clone (clone 1) was chosen for subsequent experiments.

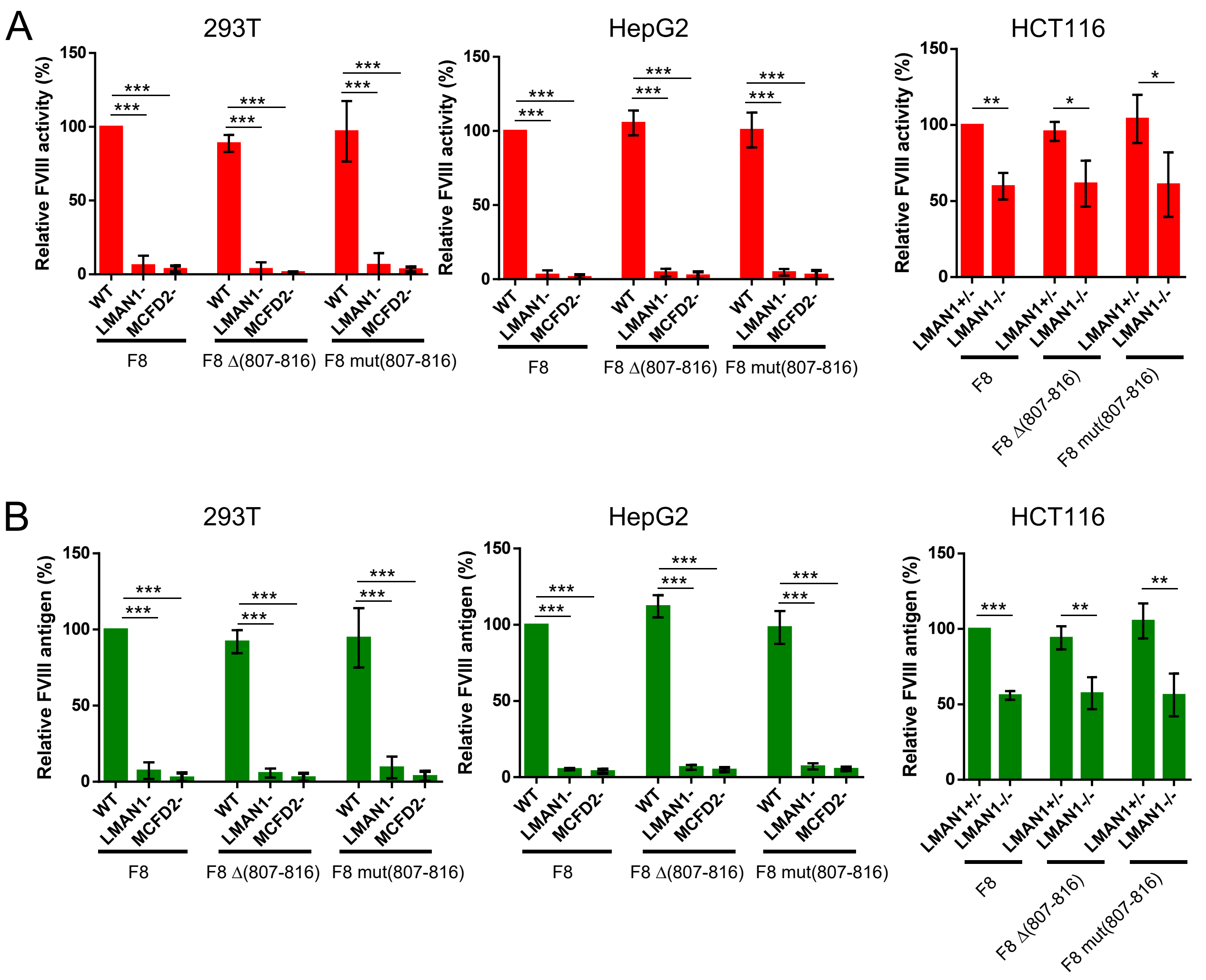


**Figure S2. Secretion levels of F8Δ (807-816) and F8mut (807-816) in *LMAN1* or *MCFD2* KO cell lines.** F8Δ (807-816) and F8mut (807-816) plasmids were transfected into different cell lines, and conditioned media were collected 48 h post transfection. FVIII activity and antigen levels in conditioned media were measured and plotted as percentages of WT cell levels. Data presented are means of 3 independent experiments, and the error bars represent standard deviations. *P<0.05, **P<0.01, ***P<0.001.

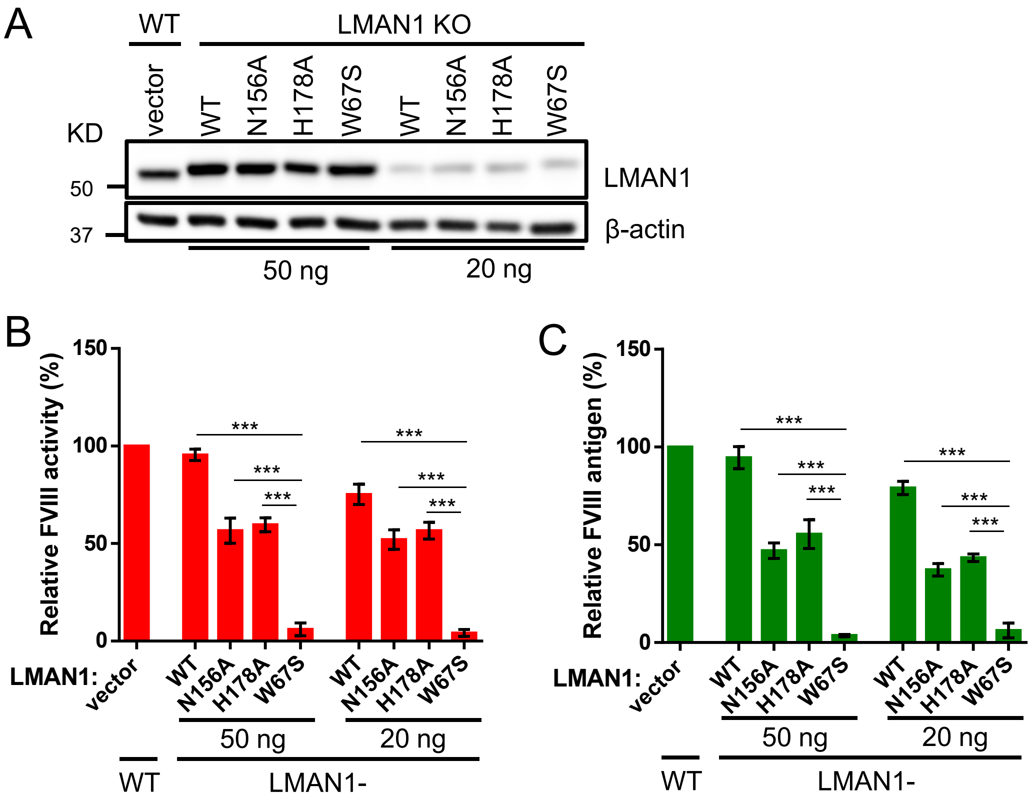


**Figure S3. Rescue of FVIII secretion in LMAN1 KO cells by LMAN1 variants with different plasmid amount.** (A) FVIII was co-transfected with the indicated LMAN1 variants into 293T$^{LMAN1-}$ cells in two doses (50 ng and 20 ng). LMAN1 expression levels were detected by immunoblotting. FVIII activity (B) and antigen (C) levels in conditioned media were measured 48 h post transfection and plotted as percentages of WT cell levels. FVIII activity and antigen data presented are means of 3 independent experiments, and the error bars represent standard deviations. ***P<0.001.

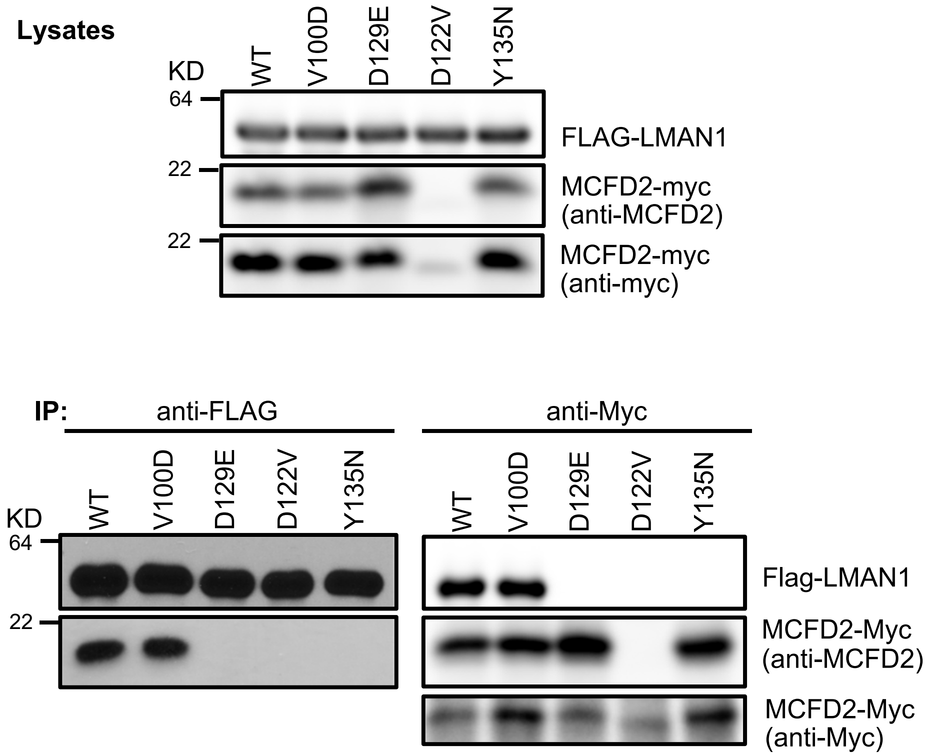


**Figure S4 Co-IP of MCFD2 variants with LMAN1**. 293T cells were co-transfected with Myc-tagged WT and MCFD2 mutants and a FLAG-tagged LMAN1. Cell lysates were immunoprecipitated with anti-Myc for MCFD2 and anti-FLAG for LMAN1 and detected by immunoblotting.

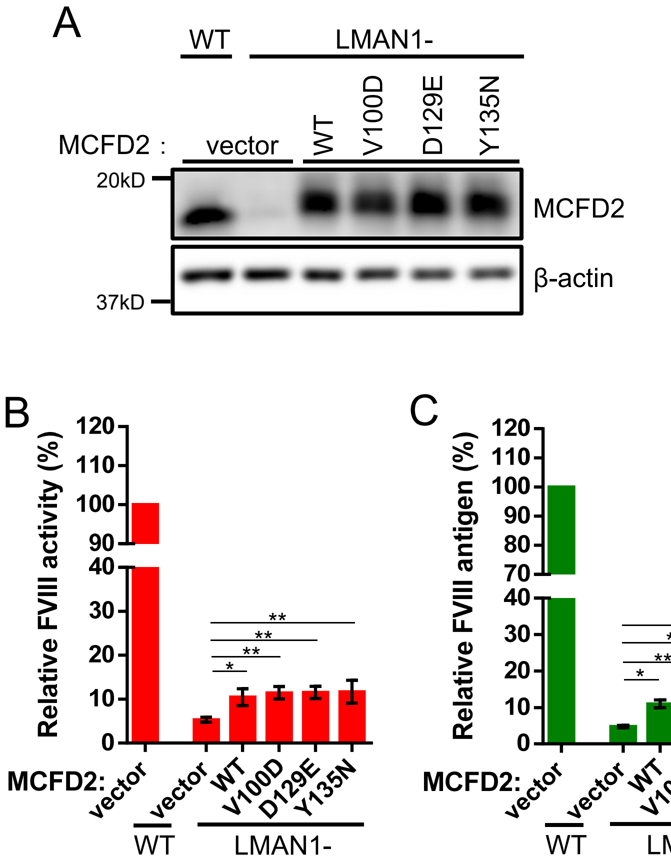


**Figure S5. FVIII secretion in LMAN1 KO cells stably expressing MCFD2 variants.** (A) WT MCFD2 and the indicated MCFD2 variants were stably expressed in 293T$^{LMAN1-}$ cells. After transfection of a FVIII expression construct, MCFD2 expression levels were compared with vector-transduced WT 293T and 293T$^{LMAN1-}$cells by immunoblotting. FVIII activity (B) and antigen (C) levels in conditioned media were measured and plotted as percentages of WT cell levels. FVIII level data presented in this figure are means of 3 independent experiments, and the error bars represent standard deviations. *P<0.05, **P<0.01

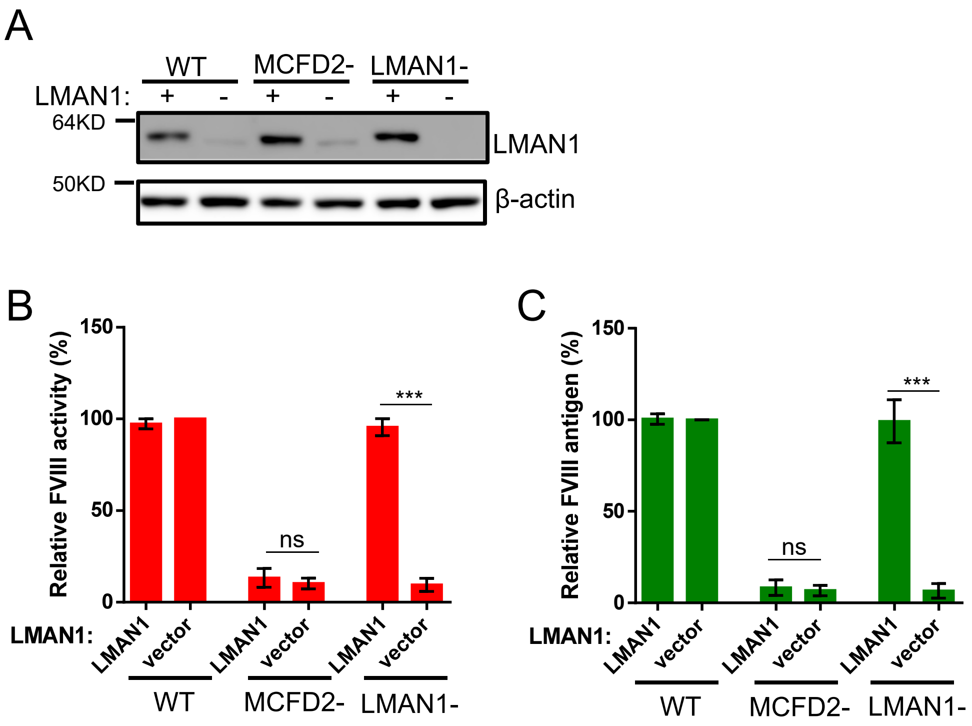


**Figure S6. Overexpression of LMAN1 cannot rescue FVIII secretion in MCFD2 KO cells.** (A) A FVIII expression construct and vector expressing WT LMAN1 were co-transfected into WT 293T, $293T^{LMAN1-}$ and $293T^{MCFD2-}$ cells. LMAN1 expression levels were detected by immunoblotting. FVIII activity (B) and antigen (C) levels in conditioned media were measured 48 h post transfection and plotted as percentages of WT cell levels. FVIII activity and antigen data presented are means of 3 independent experiments, and the error bars represent standard deviations. ***P<0.001. ns, not significant.

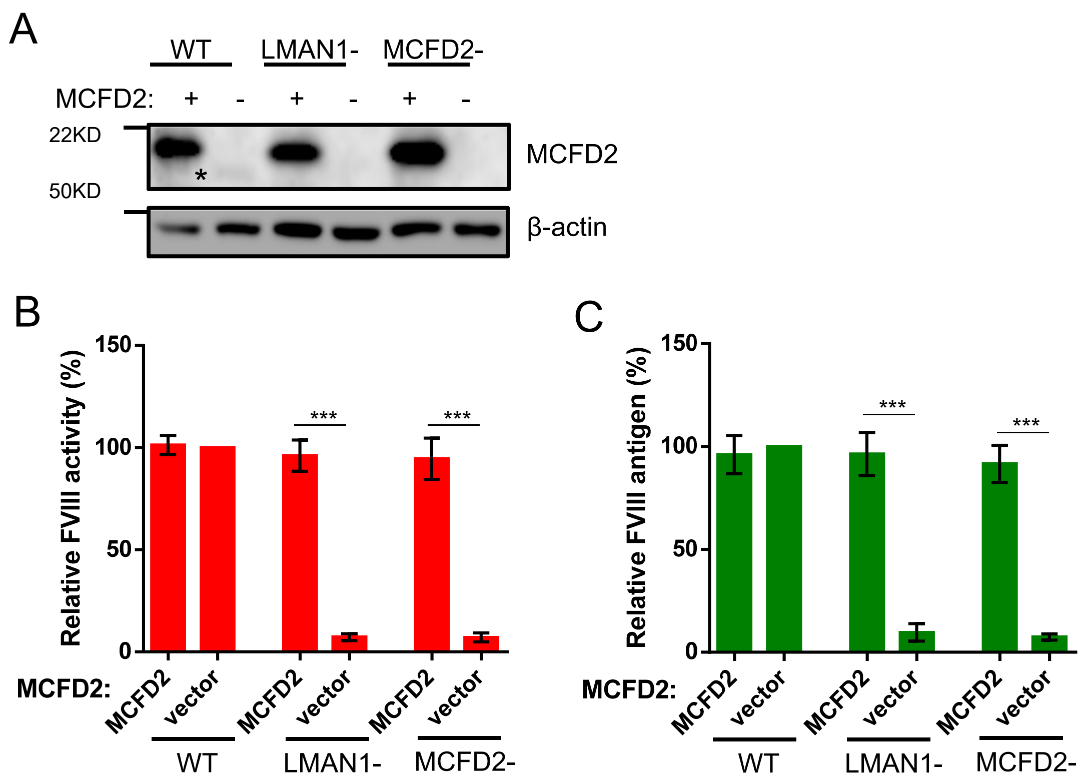


**Figure S7. Overexpression of MCFD2 rescues FVIIIΔ.807-816) secretion in LMAN1 KO cells.** (A) F8Δ(807-816) and MCFD2 expression plasmids were co-transfected into WT 293T, $293T^{LMAN1-}$ and $293T^{MCFD2-}$ cells. MCFD2 expression levels were detected by immunoblotting. FVIII activity (B) and antigen (C) levels in conditioned media were measured 48 h post transfection and plotted as percentages of WT cell levels. FVIII activity and antigen data presented are means of 3 independent experiments, and the error bars represent standard deviations. ***P<0.001.
